# The rhythmic bimodal sensory stimulation in synchronous manner entrains the network oscillation in basolateral amygdala

**DOI:** 10.1101/2025.08.22.670247

**Authors:** Miki Hashizume, Rina Ito, Ayumi Hirao, Yasushi Hojo, Gen Murakami, Takayuki Murakoshi, Naonori Uozumi

**Affiliations:** Department of Biochemistry, Faculty of Medicine, Saitama Medical University; Department of Liberal Arts, Faculty of Medicine, Saitama Medical University

## Abstract

The state of neural oscillation is important for various brain functions. In the basolateral nucleus of amygdala (BA), the oscillation frequency is accelerated in retrieval of conditioned fear memory. The amygdala receives sensory inputs from associated cortex and thalamus. Therefore, we tried to apply the bimodal sensory stimulation at slow frequency (5 Hz, functional frequency in behavioral context) for the entrainment of the BA oscillation. Young adult rats (P24-30) were stimulated by LED illuminator and acoustic speaker at 1 or 5 Hz for 1 hour. Immediately after the stimulus was finished, BA slices were prepared and whole-cell recording was applied to projection neuron. The slow (0.5-2 Hz) rhythmic IPSCs obtained from the pyramidal neuron was accelerated at ∼4 Hz by synchronous opto-acoustic stimulation at 5 Hz. However, the frequency of the neuron at the later recording did not change in the same slice, suggesting that this induced entrainment is transient and reversible phenomenon. As a result, the power distribution was shifted from 0.1-2 to 2-6 Hz by synchronous bimodal 5 Hz stimulation. The regularity of the interval between IPSCs, quantified by rhythm index and the concentration of power around peak frequency in the power spectrum, was not changed by rhythmic sensory stimulation. These results suggest that synchronous bimodal sensory stimuli control the neuronal oscillation frequency by applying with rhythmicity.

## Introduction

Neural oscillation is observed in various brain regions at variable frequencies, and is involved in physiological and pathological condition. Theta oscillation (4-12 Hz) is observed in the hippocampus and involved in functions including the spacial memory (Boyce et al., 2016; Ognjanovski et al., 2017; Ognjanovski et al., 2018). Gamma (30-80 Hz) oscillation in the hippocampus and the olfactory bulb was impaired in the Alzheimer’s disease (Iaccarino et al., 2016) and depression (Li et al., 2023), respectively. Auditory or opto-acoustic stimulation at 40 Hz was applied to Alzheimer’s animal models for restoration of behavioral and molecular impairment (Martorell et al., 2019; Murdock et al., 2024). These rhythmic sensory inputs achieved the entrainment of 40 Hz oscillation in local field potential in the hippocampus of intact and diseased animals, and reduced the amyloid level and improved spacial memory in model animals (Adaikkan et al., 2019). These findings suggested that rhythmic noninvasive stimulation induced the entrainment of brain oscillation and restored its function.

Basolateral amygdaloid complex (BLA) is essential for emotional behavior, especially for cue-dependent and contextual-dependent fear conditioning. The BLA receives multiple sensory signals from thalamus and sensory cortices, so that neutral tone or light signals associated with electrical shock can produce aversive emotion in the fear-conditioned animals. It is also reported that projection neurons of basolateral nucleus (BA, a ventral part of BLA) showed slow rhythmic activities and that they burst-fired at a low rate (∼1 Hz) preferentially during slow-wave sleep (Paré and Gaudreau, 1996). Similarly, in vitro electrophysiological studies revealed that BA projection neurons receive slow (0.5-2 Hz) rhythmic inhibitory inputs driven by the local neural circuit consisting of GABAergic and glutamatergic transmissions (Popescu and Paré, 2011). Our present study aimed to investigate whether the frequency of brain oscillation is modified by rhythmic sensory stimulation. Especially, we tried to establish the entrainment of BA activity into theta (4-10 Hz) oscillation, which has been often observed in BLA when the conditioned animal experienced fear-reminding situation (Seidenbecher et al., 2003; Karalis et al., 2016). Therefore, it is eventually expected that entrained theta oscillation in BLA enhances learning process of the fear memory.

Here we reported that the entrainment of BA oscillation was achieved only when synchronous bimodal sensory stimulation was applied to the rats, and the entrained frequency reversed gradually. The power distribution was also shifted to faster frequency with keeping the regularity of occurrence of inputs.

### Experimental procedures

#### Animals and opto-acoustic stimulation

Wistar rats (male = 33, female = 22) at P24-30 were purchased from Tokyo Laboratory Animals Science (Tokyo, Japan). They were housed with their mothers in a 12/12 h light–dark cycle (light: 7:00–19:00, dark: 19:00–7:00). Then, they were separated individually in their home cages immediately before the experiments. Thereafter, they were used for the following stimulation protocols (Fig 1A). All animal experiments were performed in accordance with the experimental protocol approved by the Institutional Animal Care and Use Committee of Saitama Medical University (approval ID #:3753).

**Figure 1.**
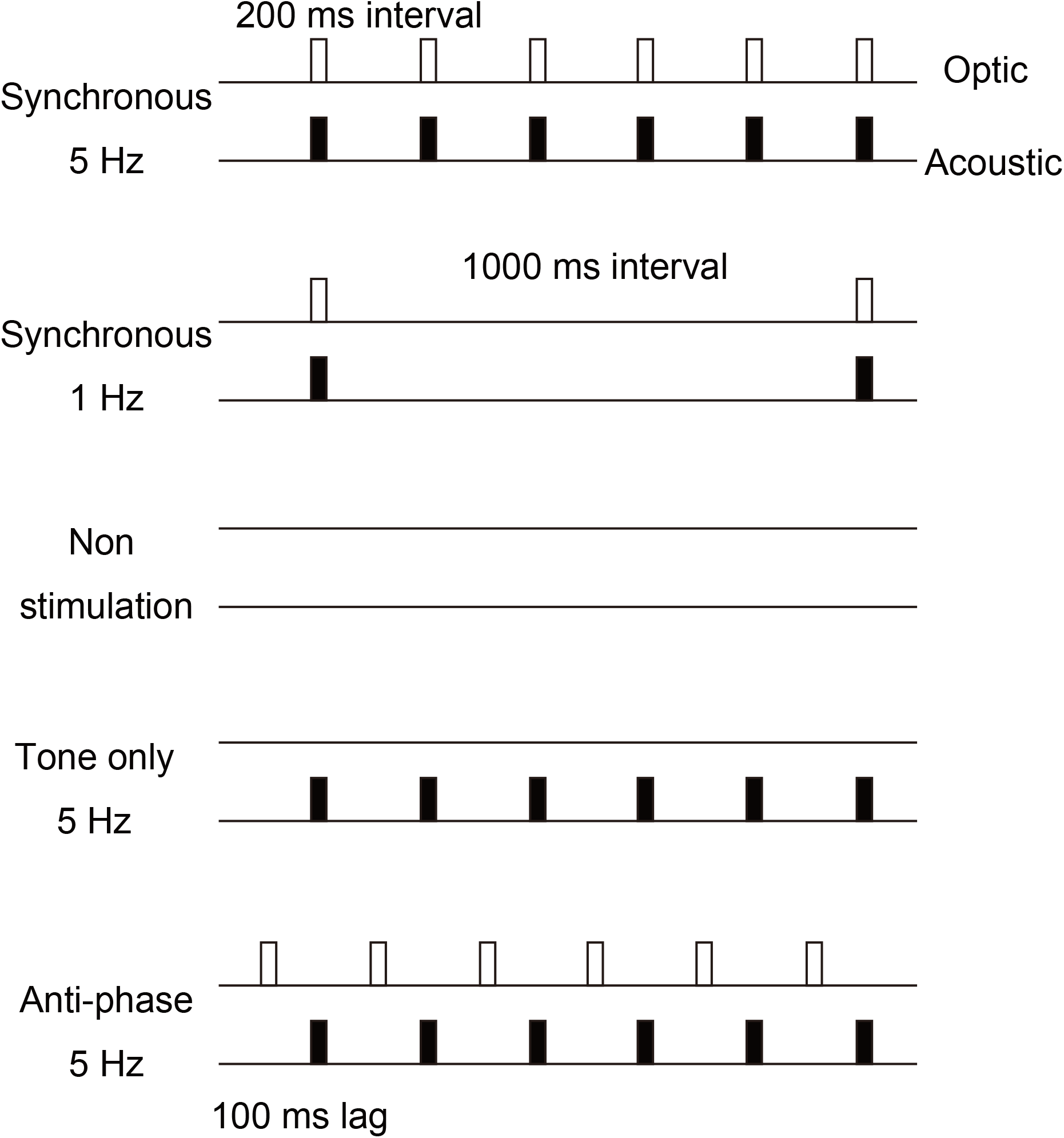
The sensory stimulation protocol Optic (white) and acoustic (black) stimulus were delivered synchronously with 200 ms or 1000 ms interval in Synchronous 5 Hz (Syn5) or Synchronous 1 Hz group (Syn1), respectively. In Anti-phase 5 Hz group (Anti5), optic stimulus was delivered at 5 Hz frequency followed by 5 Hz acoustic stimulus with 100 ms lag time.

Opto-acoustic stimulation was performed between 11:00-12:00. The rats were located in the cage (32 × 21 × 12 cm). The light (100-1000 Lux, 25 ms) and tone (10 kHz, 50 dB, 25 ms) were provided from LED illuminator and speaker, respectively. During one hour, tone was delivered with light synchronously at 5 Hz (Synchronous 5 Hz, Syn5) or 1 Hz (Synchronous 1 Hz, Syn1), or in anti-phase at 5 Hz (Anti-phase 5 Hz, Anti5), or without light at 5 Hz (Tone only 5 Hz, Tone5; see Fig 1).

#### Slice preparation

Rats were anesthetized with isoflurane immediately after the opto-acoustic stimulation. Before slicing, the rats were transcardially perfused with cooled cutting solution containing the following (in mM): 120 choline chloride, 3 KCl, 28 NaHCO_3_, 1.25 NaH_2_PO_4_, 22 glucose, and 8 MgCl_2_. The solution was oxygenated with a mixture of 95% O_2_ and 5% CO_2_. After decapitation, the brains were dissected from the rats and soaked in ice-cold cutting solution. Coronal brain slices (400 μm thick), including the amygdala, were sectioned using a microslicer (Leica VT1000S, Germany) and incubated for at least 1 h in an oxygenated artificial cerebrospinal fluid (ACSF) containing the following (in mM):120 NaCl, 3 KCl, 26 NaHCO_3_, 1.25 NaH_2_PO_4_, 15 glucose, 2.5 CaCl_2_ and 1.3 MgCl_2_. Then, the slices were transferred onto the recording chamber perfused with warm (30–32 °C) oxygenated ACSF.

#### Electrophysiology

Whole-cell patch-clamp recording was performed using a micropipette (5–9 MΩ; WPI, 1B150F-4, USA) filled with an internal solution containing the following (in mM):150 K-methanesulfonate, 5 KCl, 0.1 K-EGTA, 5 Na-HEPES, 3 Mg-ATP and 0.4 Na-GTP (pH 7.4). Whole-cell recording data were obtained from 1–4 neurons per rat. To record inhibitory postsynaptic currents/potentials (IPSCs/IPSPs) from the projection neurons, the recording neuron was identified as a pyramidal neuron in the BA by its pyramidal appearance, evoked firing pattern with spike adaptation, and relatively small fast afterhyperpolarization (fAHP). IPSCs in projection the neurons were recorded with a voltage-clamp configuration at a holding potential of -40 mV using Axopatch 200 B (Axon Instruments). The recording latency was defined as the time at which the IPSC recording was started. Data were acquired at 5 kHz and filtered at 1 kHz using a Digidata 1322 A (Axon Instruments).

#### Data analysis

All electrophysiological data were analyzed using Clampfit 10 (Molecular Devices) and Igor Pro (Wavemetrics). Power spectral analysis was conducted on the IPSC traces for 120 s using a fast Fourier transformation (FFT). The power spectral density (PSD) between 0.1 and 10 Hz was summed to calculate the power, which was used to evaluate the oscillation level. To confirm the acceleration of the oscillation, the power within 2-6 Hz and within 0.1-2 Hz were summed, and both were normalized by the summed power within 0.1-10 Hz.

Furthermore, to analyze the rhythmicity of the activities, autocorrelation analysis was performed using IPSC traces, and the rhythm index (RI) was calculated as reported previously (Lang et al., 1999; Hashizume et al., 2019). Briefly, the baseline of the autocorrelation histogram was defined as the average value from 0.1-5 s. The set of peaks and troughs was determined when the value was larger or smaller than the baseline by 1 S.D. Thereafter, the RI was determined as the maximal difference between the peaks and troughs, except for time zero. Moreover, a zero value was applied when no peaks or troughs over 1 S.D. of the signals were detected.

To quantify the fidelity of oscillatory IPSCs, the concentration of power around the peak frequency (PF) was calculated from the PSD. The proximity of PF was defined as the region which was compartmentalized between the smallest and the largest abscissas of half peak. The area of PF proximity was divided by total (0.1-10 Hz) area of power. The average height of PF proximal region, calculated by dividing the area by the width, was defined as “spectral prominence (SP)”.

#### Statistical analysis

Nonparametric data were plotted using a whisker and box plot, and compared using the Kruskal-Wallis test and post hoc Steel Dwass test. Parametric data were plotted using a bar chart, and the difference was analyzed using the one-way ANOVA test and post hoc Tukey test. The correlation between two parameters was tested by Pearson’s test. All statistical analyses were performed using the R software (https://cran.r-project.org/). The criterion for statistical significance was set at p < 0.05.

## Results

### Synchronous 5 Hz stimulation accelerates the slow inhibitory oscillation in BA projection neurons

In the BA, rhythmic IPSCs are observed in projection neuron at 0.5-2 Hz (Rainnie, 1999; Rainnie et al., 2004; Ohshiro et al., 2011; Popescu and Paré, 2011). We tried to entrain this inhibitory oscillation to applied frequency. The frequency of inhibitory oscillation in BA projection neuron was around 4 Hz after synchronous opto-acoustic stimulation at 5 Hz was applied to the rat for 1 hour (Fig 2A, upper). It was noted that another neuron in the same slice had slower frequency (around 2 Hz) in later recording (Fig 2A, lower). The latter was similar to the data in other experimental groups (Non-stimulation, Fig 2B). This tendency was confirmed by plotting the peak frequency against the recording latency (Fig 3A). The PF from synchronous 5 Hz stimulated group (Syn5) was decreased as the recording latency was later (n = 50, r = -0.31, p = 0.028). On the other hand, anti-phase 5 Hz (Anti5; n= 20, r = -0.07, p > 0.1) and tone only 5 Hz (Tone5; n= 10, r = 0.13, p > 0.1) stimulated groups showed no significant correlation, same as the results of synchronous 1 Hz (Syn1; n= 22, r = -0.12, p > 0.1) and non stimulation groups (Non; n = 22, r = -0.06, p > 0.1). Moreover, the values of PF in synchronous 5 Hz stimulated group was significantly larger than that in the other experimental groups (Fig 3B, X^2^ = 28.4, p = 0.0000104). Other stimulus protocols did not change the frequency from original slow (0.5-2 Hz) oscillatory activity. Besides the PF change, the shift of power distribution was analyzed by calculating the area under curve of PSD. The range we applied were 0.1-2 Hz and 2-6 Hz, since previous research revealed that 4-Hz oscillation (2-6 Hz) in BLA was strongly correlated with fear memory retrieval (Karalis et al., 2016). The power was concentrated within faster frequency range by synchronous 5 Hz stimulation, while the power within original frequency range was decreased (Fig 4A). These results suggest that synchronous bimodal sensory stimulation can entrain the oscillation at designed frequency, and this entrainment is reversible phenomenon.

**Figure 2.**
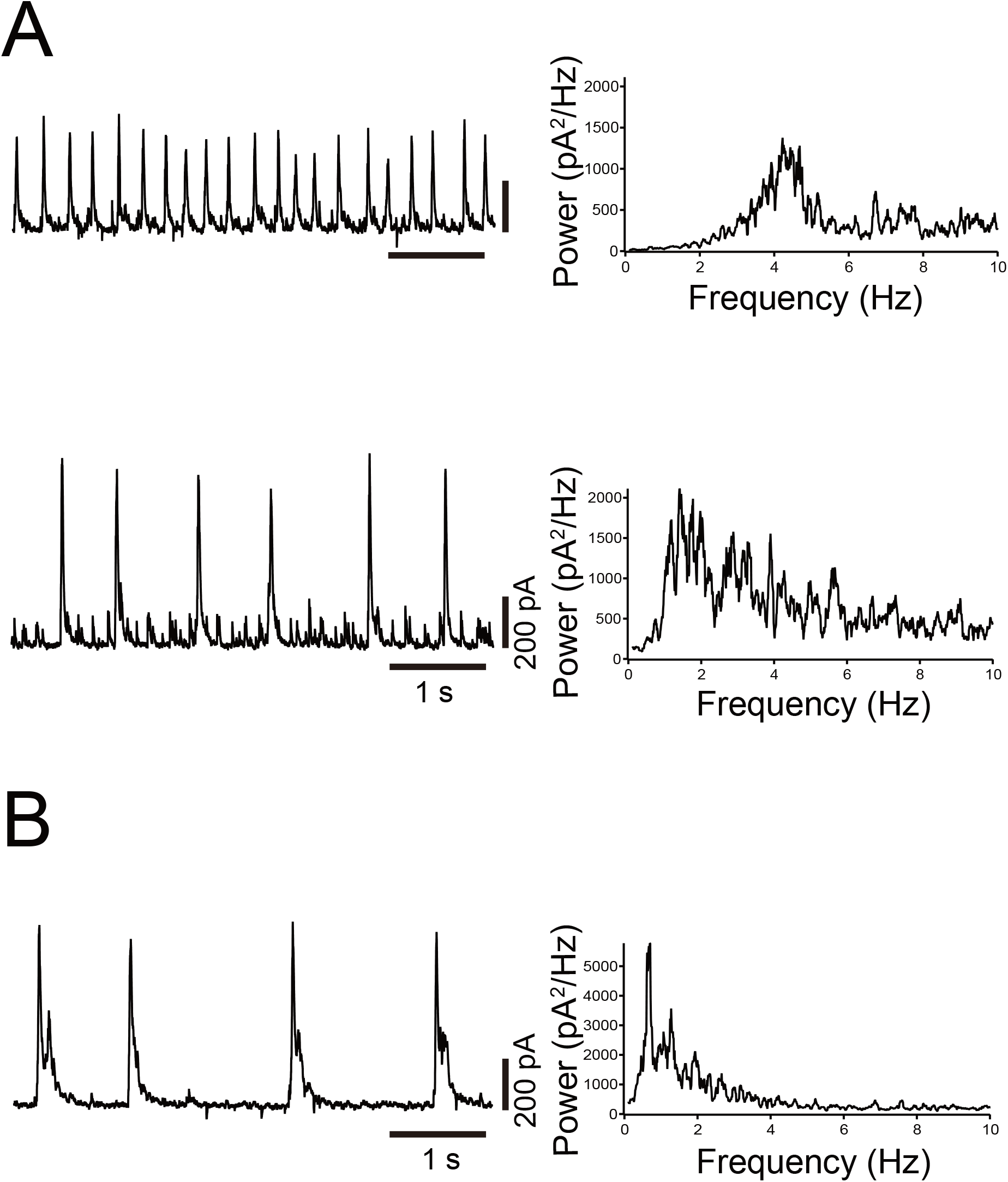
Oscillatory IPSCs in BA projection neuron were entrained by rhythmic sensory stimuli (A) Oscillatory IPSCs recorded at -40 mV was accelerated by synchronous 5 Hz tone-light stimulation in BA projection neuron at early recording (upper). Power spectral analysis showed that the peak frequency was shifted at ∼4 Hz. Another neuron in the same slice in A showed the slower oscillation at later recording (lower). The peak frequency was within original value (0.5-2 Hz). (B) Oscillatory IPSCs was obtained from the BA neuron in non-stimulated rat. The peak frequency was within original value (0.5-2 Hz).

**Figure 3.**
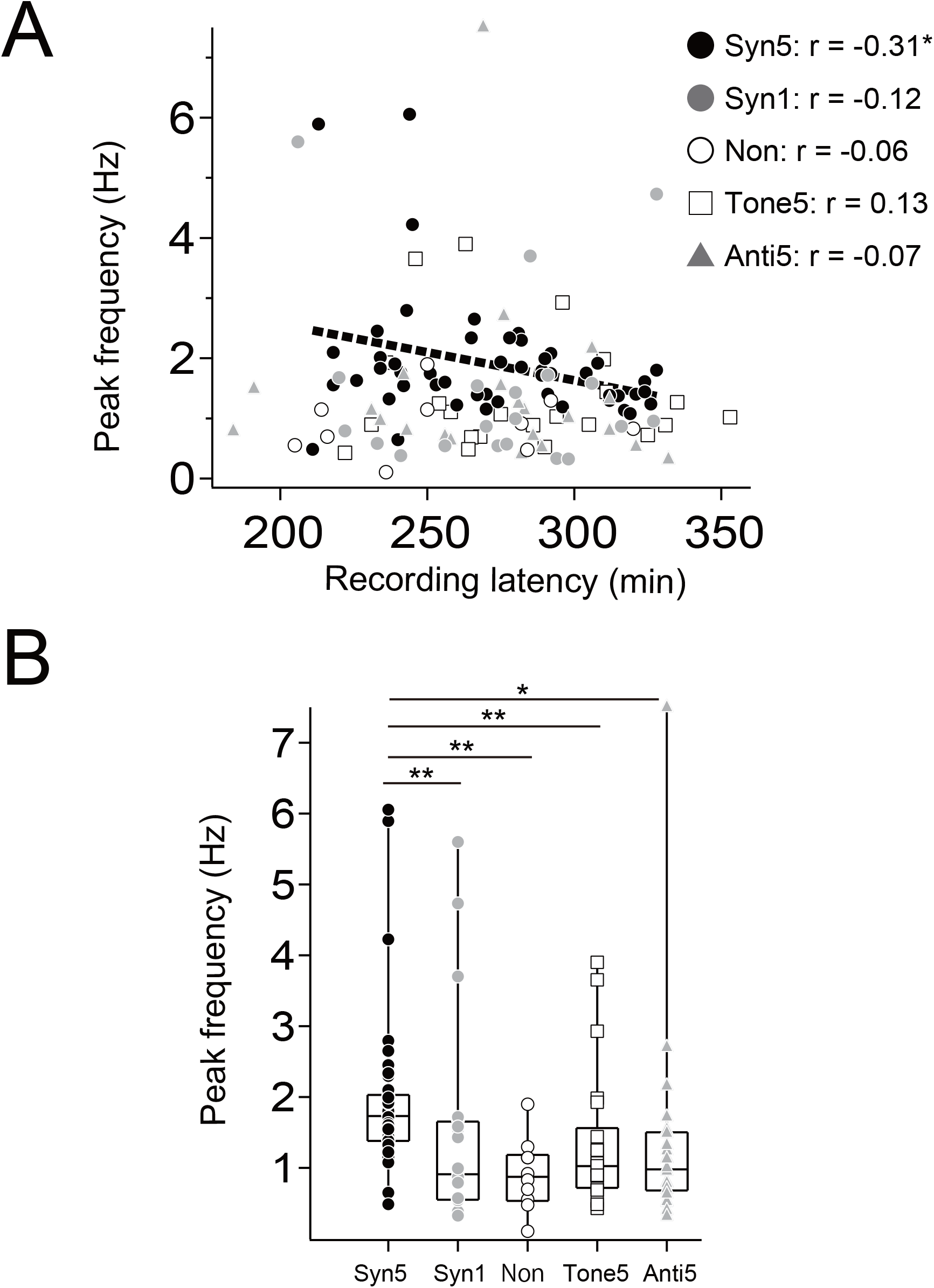
The peak frequency entrained by synchronous oscillatory stimuli was returned gradually (A) The peak frequency was plotted against recording latency, which was defined as the time at which the IPSC recording was started (0 min was the time when the rat was sacrificed by isoflurane). The data of Synchronous 5 Hz (black circle) had significant negative correlation (r = 0.31, *p < 0.05). (B) peak frequency of Synchronous 5 Hz was only significantly faster than those of other four experimental groups (**p < 0.01, *p < 0.05).

**Figure 4.**
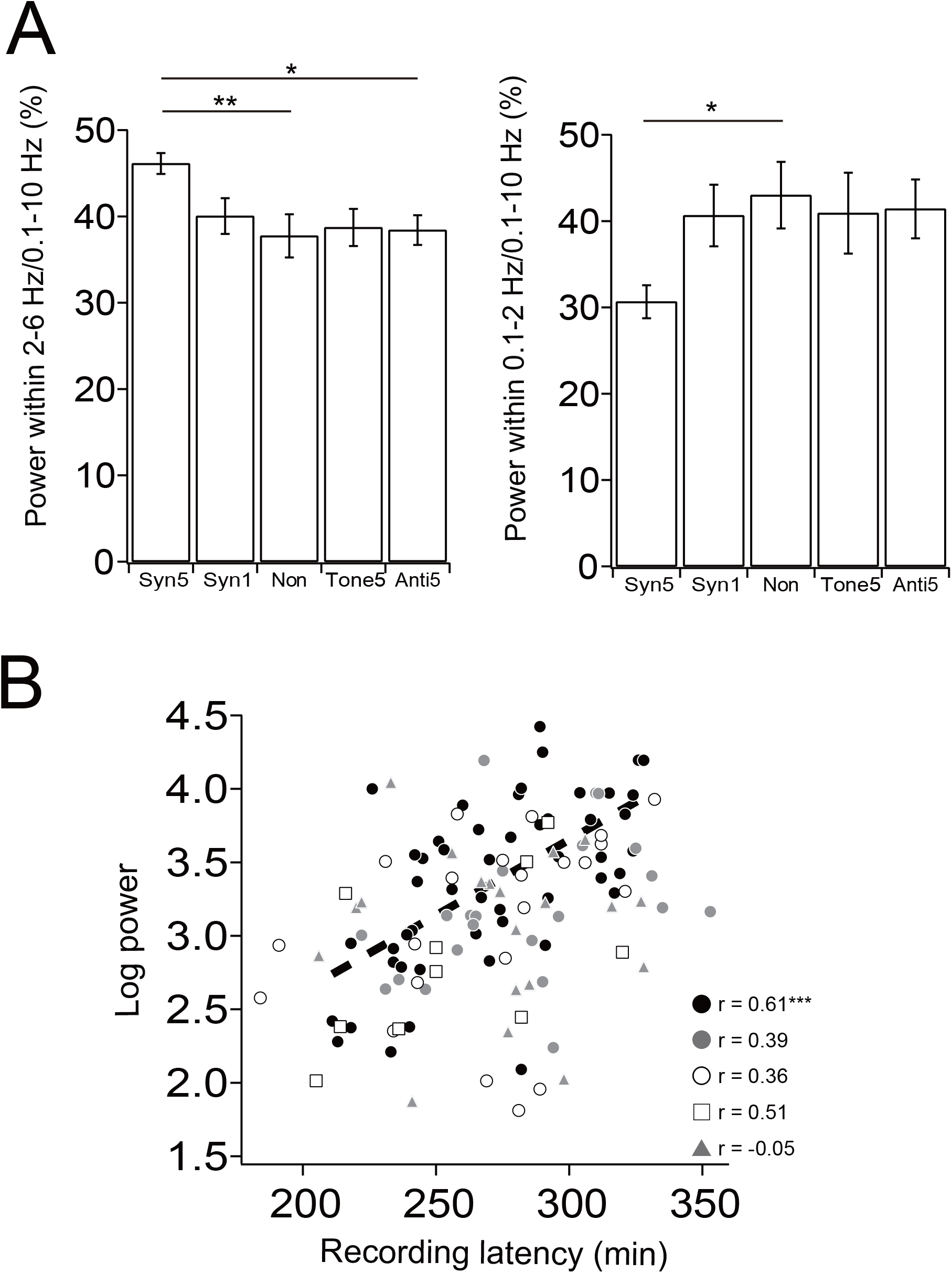
The oscillation power (A) Summed oscillation power within low range. Oscillation power within 2-6 Hz were plotted (left), and power within 0.1-2 Hz were plotted (right). (B) The log power was plotted against recording latency. The data of Synchronous 5 Hz (black circle) had significant positive correlation (r = 0.61, ***p < 0.001).

There was no significant different between the oscillation powers in sensory stimulated BA and that in non-stimulated BA (Table 1). However, there was significant positive correlation between the power in synchronous 5 Hz and recording latency (Fig 4B), suggesting the synchronous bimodal sensory stimulation affected the oscillation power besides the PF.

**Table 1.** Summerized oscillation parameters in all experimental groups. The median and 95% confidential interval were displayed. Top row: The integrated oscillation power within 0.1-10 Hz. Bottom row: The rhythm index (RI) was calculated from autocorrelation analysis. There were no significant difference among all experimental groups.

|  | Syn5 | Syn1 | Non | Tone5 | Anti5 |
| --- | --- | --- | --- | --- | --- |
| Log power | 3.47<br>(3.20-3.53) | 3.14<br>(2.97-3.39) | 3.35<br>(2.82-3.39) | 2.82<br>(2.44-3.23) | 3.20<br>(2.79-3.30) |
| Rhythm<br>index | 0.19<br>(0.16-0.22) | 0.12<br>(0.10-0.15) | 0.15<br>(0.10-0.20) | 0.12<br>(0.03-0.16) | 0.10<br>(0.09-0.19) |

### The analysis of rhythmic IPSC fidelity

External sensory stimulation in present study was completely periodic (200 or 1000 milliseconds interval). Therefore, the oscillatory activity in BA might become more periodic by the stimulation. To confirm this possibility, the fidelity of the oscillatory event (IPSC) was quantified by calculating the spectral prominence (SP; see Methods), the ratio of concentration of power around PF. PSD was made from IPSC trace and SP was calculated using half peak value (Fig 5A). SP was higher when the peak of spectrum was narrow and prominent (Fig 5A, top), and was lower when multiple peaks (or a peak and shoulders) was emerged (Fig 5A, bottom). There was no significant difference in SP among five experimental groups (Fig 5B).

**Figure 5.**
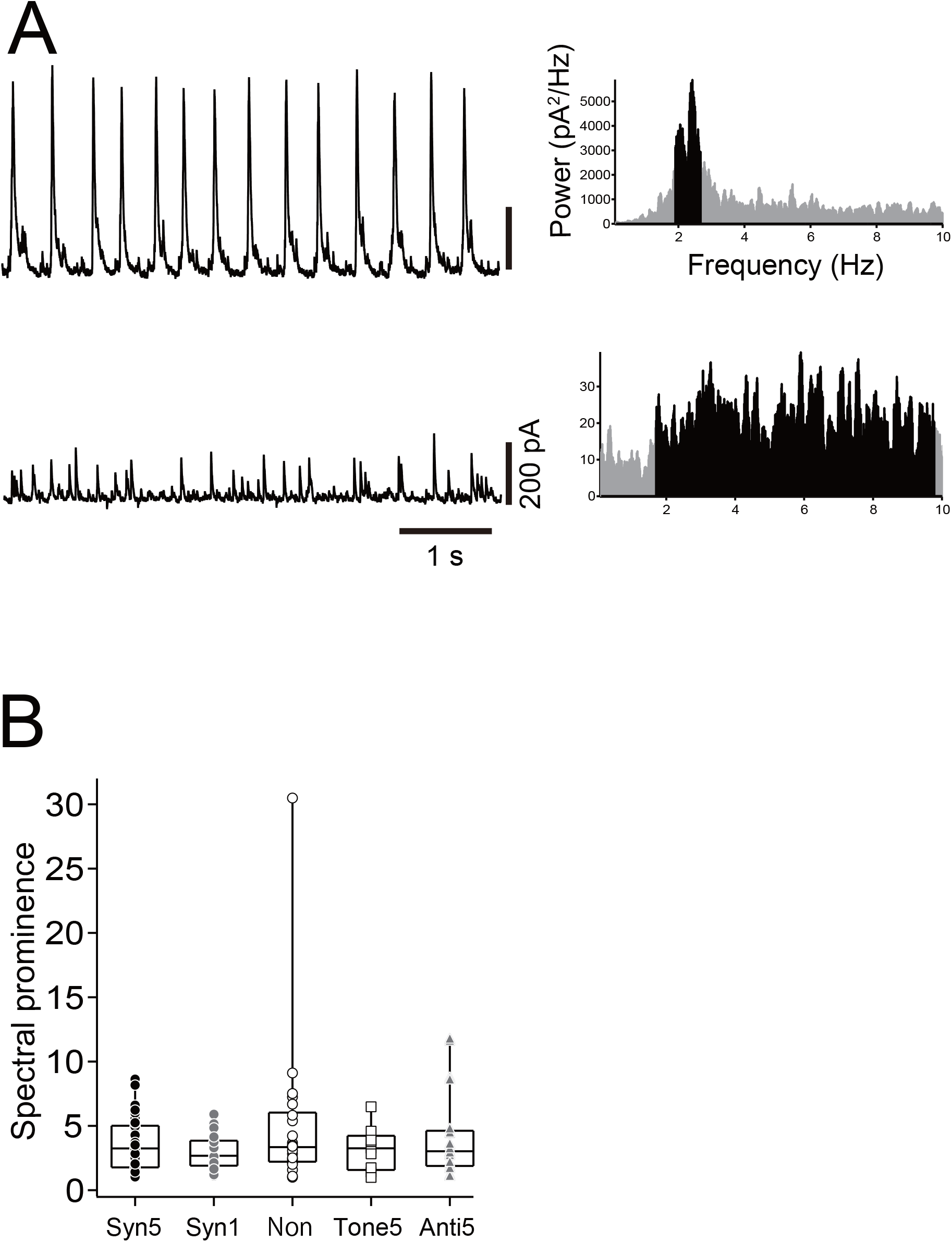
The spectral prominence (SP) was not changed by external sensory oscillatory stimuli (A) Power spectral analysis (right), which was made from IPSC trace (left), was used for calculation of SP. Around peak (black area) was defined by the first- and the last-emerged half peak abscissas. In upper case, the range around peak was narrower, so that the SP contributed to the area compared with the width (SP was higher). In lower case, the range around peak was wider, so that the width contributed to the area compared with the SP (SP was lower). (B) SP, calculated by dividing the black area by the width, were plotted. There was no significant difference among five experimental groups.

Similar to SP, rhythm index (RI) calculated from autocorrelogram, had no significant difference among five experimental groups (Table 1; X^2^ = 11.4, p > 0.05). The SP was plotted against RI for investigation of relationship (Fig S1). There was significant correlation in 5 Hz synchronous (r = 0.73), 5 Hz anti-phase (r = 0.76), 5 Hz tone only (r = 0.89) and no stimulation group (r = 0.85). The result suggested that RI and SP is the similar criterion about rhythmicity.

## Discussion

The results of the present study showed that rhythmic bimodal sensory (optic/acoustic) stimulation in synchronous manner (5 Hz) was able to induce entrainment of the oscillation in the basolateral nucleus of amygdala (BA) projection neurons, while the mono-sensory (acoustic) stimulation at the same frequency (5 Hz) did not induce entrainment. Earlier studies reported that the entrainment was induced by mono-sensory rhythmic stimulation in auditory thalamus or visual cortex (Herrmann, 2001; Gao et al., 2009). Although the lateral nucleus of Amygdala receives the output from the auditory thalamus and cortex, rhythmic mono-sensory input failed to be affected on BA. Moreover, the coincidence was also important for the entrainment in BA, since 5 Hz bimodal sensory stimulation in anti-phase manner failed to induce the entrainment. In cue-associated fear conditioning in BLA, sensory signal pathway associated with neutral cue is potentiated when aversive stimulus and neutral cue are coincidently applied. Similar to that, multimodal sensory inputs in synchronous manner is required for the entrainment in amygdala oscillation.

The synchronous sensory stimulation also affected the power of BA oscillation. The oscillation power was increased as the recording latency was later (Fig 4B), while there was no significant difference among five experimental groups (Table 1). Our studies revealed that the power was decreased by stressful condition and recovered slowly (Hashizume et al., 2019; Hashizume et al., 2024), suggesting that present opto-acoustic stimulation had little stressful effect on rats.

The change of peak frequency lasted for 2-3 hours after the slice was prepared and gradually reversed (Fig 2 and 3). The persistence of entrainment was reported in the auditory thalamus of anesthetized mice, in which the entrainment induced by rhythmic sound stimuli for 2 minutes was kept for ∼1 minute (Gao et al., 2009). These results suggest that long lasting plastic change (protein synthesis, synaptogenesis) is neither involved in the oscillation entrainment, and nor needed for the function. The frequency of electroencephalogram (EEG) is altered according to the sleep state, or the BLA local field potential (LFP) was changed by the animal behavior (Karalis et al., 2016). Therefore, the reversibility of oscillation must be kept for the representation of brain activity.

The oscillatory fidelity of IPSCs was not changed by rhythmic sensory stimulation. There was also no significant difference in rhythm index (RI) between stimulated and non-stimulated groups. Similar to that, the concentration of power around peak frequency (PF), quantified by spectral prominence (SP), was not altered by any types of sensory stimulation. Thus, synchronous 5 Hz stimulation increased the peak frequency of power (Fig 3B), induced the power distribution shift from 0.1-2 Hz to 2-6 Hz (Fig 4A), and did not decreased the regularity of IPSC occurrence (Fig 5B). In optogenetic study, direct rhythmic stimulation to cortical neurons induced the entrainment of LFP oscillation in anesthetized rats (Kuki et al., 2013). Although the entrainment could be induced at various frequency, the original PF in motor cortex (about 0.8 Hz) was kept, so that there were one peak at ∼0.8 Hz and the shoulder at the entrained frequency in the power spectrum. Our results in amygdala suggest that the BA oscillation is altered with only single peak frequency by the rhythmic stimulation.

Rhythmic sensory stimulus in noninvasive manner was applied to various neuronal diseases (Adaikkan et al., 2019; Martorell et al., 2019; Kim et al., 2024). In disease model animal, gamma (40 Hz) entrainment affected glial cells and recovered behavioral disorders. This effect on glial cell was thought to be induced by neuropeptide which was secreted by high frequency stimulus. However, these neuropeptides failed to secret by low frequency (<10 Hz) stimulation (Consolo et al., 1994; Murdock et al., 2024). Thus, high frequency noninvasive stimulation (and the entrained brain activity at high frequency) could be used for neuronal or glial control via the secretion of neuropeptide. On the other hand, our present study is not involved in neuropeptide secretion since the frequency of rhythmic sensory stimulus was low range (5 Hz). The shift to theta oscillation in the amygdala is important for the correlation with the prefrontal cortex or the hippocampus (Popa et al., 2010; Karalis et al., 2016). The rhythmic sensory stimulation is expected to induce the correlation among prefrontal and limbic regions.

## Figure legends

**Supplemental figure 1.**
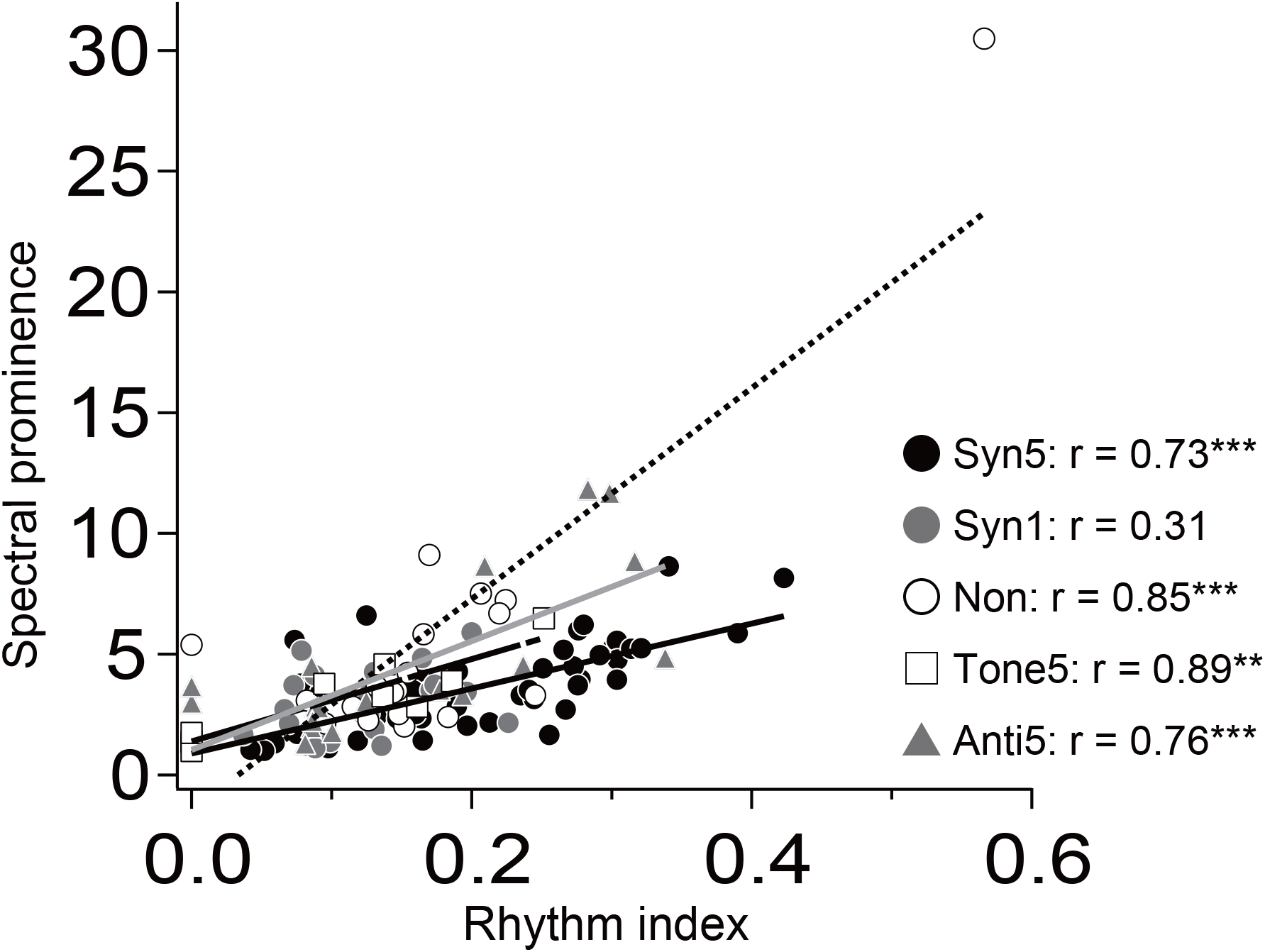
The relationship between spectral prominence (SP) and rhythm index (RI) SP was plotted against RI. Significant positive correlation were observed in Synchronous 5 Hz (black circle; r = 0.73, ***p < 0.001), No stimulation (white circle; r = 0.85, ***p < 0.001), Tone-only 5 Hz (square; r = 0.89, **p < 0.01) and Anti-phase 5 Hz groups (gray triangle; r = 0.76, ***p < 0.001). There was no significant difference in Synchronous 1 (gray circle; r = 0.31, p > 0.1).

## References

Adaikkan C, Middleton SJ, Marco A, Pao PC, Mathys H, Kim DN, Gao F, Young JZ, Suk HJ, Boyden ES, McHugh TJ, Tsai LH (2019) Gamma Entrainment Binds Higher-Order Brain Regions and Offers Neuroprotection. Neuron 102:929–943 e928.

Boyce R, Glasgow SD, Williams S, Adamantidis A (2016) Causal evidence for the role of REM sleep theta rhythm in contextual memory consolidation. Science 352:812–816.

Consolo S, Baldi G, Russi G, Civenni G, Bartfai T, Vezzani A (1994) Impulse flow dependency of galanin release in vivo in the rat ventral hippocampus. Proc Natl Acad Sci U S A 91:8047–8051.

Gao L, Meng X, Ye C, Zhang H, Liu C, Dan Y, Poo MM, He J, Zhang X (2009) Entrainment of slow oscillations of auditory thalamic neurons by repetitive sound stimuli. J Neurosci 29:6013–6021.

Hashizume M, Ito R, Hojo Y, Yanagawa Y, Murakoshi T (2019) Acute sleep deprivation reduces oscillatory network inhibition in the young rat basolateral amygdala. Neuroscience 401:73–83.

Hashizume M, Ito R, Suge R, Hojo Y, Murakami G, Murakoshi T (2024) Correlation Between Cued Fear Memory Retrieval and Oscillatory Network Inhibition in the Amygdala Is Disrupted by Acute REM Sleep Deprivation. Neuroscience 536:12–20.

Herrmann CS (2001) Human EEG responses to 1-100 Hz flicker: resonance phenomena in visual cortex and their potential correlation to cognitive phenomena. Exp Brain Res 137:346–353.

Iaccarino HF, Singer AC, Martorell AJ, Rudenko A, Gao F, Gillingham TZ, Mathys H, Seo J, Kritskiy O, Abdurrob F, Adaikkan C, Canter RG, Rueda R, Brown EN, Boyden ES, Tsai LH (2016) Gamma frequency entrainment attenuates amyloid load and modifies microglia. Nature 540:230–235.

Karalis N, Dejean C, Chaudun F, Khoder S, Rozeske RR, Wurtz H, Bagur S, Benchenane K, Sirota A, Courtin J, Herry C (2016) 4-Hz oscillations synchronize prefrontal-amygdala circuits during fear behavior. Nat Neurosci 19:605–612.

Kim T, James BT, Kahn MC, Blanco-Duque C, Abdurrob F, Islam MR, Lavoie NS, Kellis M, Tsai LH (2024) Gamma entrainment using audiovisual stimuli alleviates chemobrain pathology and cognitive impairment induced by chemotherapy in mice. Sci Transl Med 16:eadf4601.

Kuki T, Ohshiro T, Ito S, Ji ZG, Fukazawa Y, Matsuzaka Y, Yawo H, Mushiake H (2013) Frequency-dependent entrainment of neocortical slow oscillation to repeated optogenetic stimulation in the anesthetized rat. Neurosci Res 75:35–45.

Lang EJ, Sugihara I, Welsh JP, Llinás R (1999) Patterns of spontaneous purkinje cell complex spike activity in the awake rat. J Neurosci 19:2728–2739.

Li Q, Takeuchi Y, Wang J, Gellert L, Barcsai L, Pedraza LK, Nagy AJ, Kozak G, Nakai S, Kato S, Kobayashi K, Ohsawa M, Horvath G, Kekesi G, Lorincz ML, Devinsky O, Buzsaki G, Berenyi A (2023) Reinstating olfactory bulb-derived limbic gamma oscillations alleviates depression-like behavioral deficits in rodents. Neuron 111:2065–2075 e2065.

Martorell AJ, Paulson AL, Suk HJ, Abdurrob F, Drummond GT, Guan W, Young JZ, Kim DN, Kritskiy O, Barker SJ, Mangena V, Prince SM, Brown EN, Chung K, Boyden ES, Singer AC, Tsai LH (2019) Multi-sensory Gamma Stimulation Ameliorates Alzheimer’s-Associated Pathology and Improves Cognition. Cell 177:256–271 e222.

Murdock MH et al. (2024) Multisensory gamma stimulation promotes glymphatic clearance of amyloid. Nature 627:149–156.

Ognjanovski N, Broussard C, Zochowski M, Aton SJ (2018) Hippocampal network oscillations rescue memory consolidation deficits caused by sleep loss. Cereb Cortex 28:3711–3723.

Ognjanovski N, Schaeffer S, Wu J, Mofakham S, Maruyama D, Zochowski M, Aton SJ (2017) Parvalbumin-expressing interneurons coordinate hippocampal network dynamics required for memory consolidation. Nat Commun 8:15039.

Ohshiro H, Kubota S, Murakoshi T (2011) Dopaminergic modulation of oscillatory network inhibition in the rat basolateral amygdala depends on initial activity state. Neuropharmacology 61:857–866.

Paré D, Gaudreau H (1996) Projection cells and interneurons of the lateral and basolateral amygdala: distinct firing patterns and differential relation to theta and delta rhythms in conscious cats. J Neurosci 16:3334–3350.

Popa D, Duvarci S, Popescu AT, Léna C, Paré D (2010) Coherent amygdalocortical theta promotes fear memory consolidation during paradoxical sleep. Proc Natl Acad Sci U S A 107:6516–6519.

Popescu AT, Paré D (2011) Synaptic interactions underlying synchronized inhibition in the basal amygdala: evidence for existence of two types of projection cells. J Neurophysiol 105:687–696.

Rainnie DG (1999) Serotonergic modulation of neurotransmission in the rat basolateral amygdala. J Neurophysiol 82:69–85.

Rainnie DG, Bergeron R, Sajdyk TJ, Patil M, Gehlert DR, Shekhar A (2004) Corticotrophin releasing factor-induced synaptic plasticity in the amygdala translates stress into emotional disorders. J Neurosci 24:3471–3479.

Seidenbecher T, Laxmi TR, Stork O, Pape HC (2003) Amygdalar and hippocampal theta rhythm synchronization during fear memory retrieval. Science 301:846–850.

